# *De Novo* Regeneration of Rete Ridges during Cetacean skin wound healing

**DOI:** 10.64898/2026.03.16.711258

**Authors:** Tzu-Yu Liu, Hao-Ven Wang, Wei-Cheng Yang, Chao-Chun Yang, Chen-Yi Su, Yun-Ting Chiou, Tsyr-Huei Chiou, Shyh-Jou Shieh, Ming-Jer Tang, Cheng-Ming Chuong, Michael W. Hughes

## Abstract

Humans are tight-skinned mammals who typically fail to regenerate large full-thickness skin wounds, instead healing with substantial scarring and concomitant loss of function. Mechanical context is a major determinant of this outcome: elevated tissue tension or stiffness promotes fibrotic repair associated with hypertrophic or keloid scarring. Accordingly, regenerative medicine research has relied on diverse animal models to understand scar development and skin regeneration. Loose-skinned mammals exhibit greater regeneration ability. Furthermore, spiny mouse skin is significantly less stiff and associated with enhanced regenerative ability. Interestingly, this skin wound stiffness can be modulated to shift healing toward more regenerative or more fibrotic trajectories. Despite of this progress, the restoration of normal skin architecture after large-full thickness injury has not been elucidated in tight-skinned mammals. Can large full-thickness wounds regenerate with minimal scarring in tight-skinned mammals? Here we show the tight-skinned mammal Fraser’s Dolphin regenerates *de novo* a complex rete ridge architecture with associated vasculature and minimal scar following large full-thickness wound healing. Counterintuitively, this skin regeneration occurs in an aqueous, high-shear stress and high-tension environment. Complete rete ridge regeneration in tight-skinned mammals has not been documented and not observed in humans except *in utero*. This unique ability to rebuild elaborate rete ridges under tension is an opportunity to uncover molecular, cellular, and tissue-level mechanisms that enable regenerative wound healing in a mechanical regime typically associated with fibrosis.

## INTRODUCTION

Humans are tight-skinned mammals where the skin organ is tightly adhered to the underlying tissues (Rittie, 2016). Large full-thickness (LFT) wounds in humans and other tight-skinned mammals heal with scar formation (Shieh and Cheng, 2015). The size of the scar depends on wound location, age of the patient, and depth of the wound (Muneoka et al, 2008). The majority of clinical therapies focus on inflammation, moisture imbalance, epithelial tongue migration and scar reduction (Beraja et al, 2025). The classical wound research model for human wounds is a tight-skinned animal model the pig (Saeed and Martins-Green, 2023). The pig wound model exhibits wound contraction and scar formation very similar to humans, but does not regenerate skin structures including ectodermal organs or rete ridges (Lin et al, 2019). To our knowledge, no tight-skinned mammal has exhibited the ability to regenerate rete ridges after LFT wounding. Rete ridges are comprised of epithelium attached to the basement membrane that invaginate into the underlying papillary dermis. There are reports arguing rete ridges contain stem cells suggesting a unique skin stem cell niche (Lavker and Sun, 1983, Webb, Li and Kaur, 2004, Lin et al, 2019).

Strikingly, multiple species of loose-skinned mammals exhibit the ability to regenerate skin including ectodermal organs (Breedis, 1954, Ito et al, 2007, Seifert, Kiama et al, 2012) an enhanced regenerative ability in part due to low mechanical force (Seifert, Kiama et al, 2012, Harn, H. I. et al, 2021, Harn, Hans I-Chen et al, 2022). Reducing low mechanical force also reduces scars in humans (Gurtner, Callaghan and Longaker, 2007, Hsu et al, 2018). Unfortunately, loose-skinned animals do not develop rete ridges, and tight-skinned mammals do not regenerate rete ridges. It would be ideal to study a tight-skinned mammal that regenerates rete ridges in order to investigate this mechanism in hopes of improving human rete ridge regeneration.

Cetaceans are tight-skinned mammals inhabiting aquatic environments and accruing specific adaptations for life in a viscous watery habitat. Intimate contact with aquatic environments induced significant changes; loss of hair follicles, the replacement of suprabasal cytokeratin 10 (K10) for cytokeratin 17 (K17), and the presence of rete ridges (Ehrlich et al, 2019, Eckhart, Ehrlich and Tschachler, 2019). These changes occurred in a relatively more viscous and higher tension environment than terrestrial environments. Cetaceans evolved from an ancestral terrestrial artiodactyl that gave rise to camels, pigs, cows, giraffes, and hippopotamus (Fordyce, 2018). Most of the extant terrestrial artiodactyls do not develop rete ridges, except for suina (pigs) and the semi-aquatic hippopotamus. Hippopotamus share a more recent common ancestor with cetaceans and form a sister group (Geisler and Uhen, 2005). Humans and cetaceans do not share a recent common ancestor suggesting rete ridge presence in both could be the result of convergent evolution. While both cetaceans and humans suffer from severe LFT wounds (Honebrink et al, 2011, Mouton and Botha, 2012, Ribereau-Gayon et al, 2017), cetaceans heal remarkably well and in certain cases regenerate (Zasloff, 2011).

Here, we characterize a cetacean LFT wound healing process. Fraser’s dolphins (*Lagenodelphis hosei*) LFT wounds heal with minimal gross tissue deficit and scar. These wounds regenerate *de novo* complex rete ridge structures exhibiting elongated branching epithelium. Fraser’s dolphin rete ridge branches are intimately associated with vascularity and this association regenerates. This remarkable ability to regenerate LFT wounds in an aqueous and relatively high force environment is contrary to the human outcome of immense scarring permitting an opportunity for regenerative medicine.

## RESULTS

### Human Rete Ridges do not Regenerate After LFT Wounding

Normal and healed human skin was collected from left-over surgical excisions and rete ridges characterized (Fig. 1) Unwounded human skin exhibited undulating epithelial indentations extending into the dermis (Fig. 1a). Immunostaining for cytokeratin 14 (K14) and 10 (K10) showed normal basal and suprabasal cell patterning (Fig. 1b-c). Trichrome stain demonstrated an interlaced network of collagen fibers in the papillary dermis (Fig. 1d). PCNA and TRP63 immunostaining showed proliferation of keratinocytes only in the basal layer and normal epithelial stratification respectively (Fig. 1e-f). Integrin alpha 6 (ITGA6) and collagen XVII (COL17) exhibited a well-formed basement membrane (Fig. 1g-h). Alkaline phosphatase (ALP) staining showed positive cells at the bottom of the rete ridge that were K10-supporting a previous report (Fig. 1i) (Lin et al, 2019). The healed human lesion showed no rete ridge structures (Fig. 1j). K14 expression pattern was increased (Fig. 1k) and K10 expression pattern was reduced (Fig. 1l) suggesting abnormal keratinocyte differentiation. Trichrome staining showed a linear arrangement of collagen significantly different from normal skin (Fig. 1m). PCNA staining detected an increase in keratinocyte proliferation compared to normal skin (Fig. 1n). TRP63 showed a normal expression pattern (Fig. 1o). ITGA6 and COL17 expression patterns were normal suggesting a proper basement membrane (Fig. 1p-q). Rete ridges were absent resulting in reduced ALP staining compared to normal skin (Fig. 1r). Normal human skin epithelium was separated from the mesenchyme and used to view the three-dimensional (3-D) structure of rete ridges in whole mount studies (Fig. 1s-v). Direct illumination channel (DIC) of unstained epithelium viewed from the ventral side showed an interconnected network of rete ridges (Fig. 1s). Hoechst, ALP, and ITGA6 staining showed an interconnected network of rete ridges with ALP+ cells near the basement membrane (Fig. 1t-v). Interestingly, the ALP staining intensity was greater in the rete ridge basal cells, supporting the paraffin section data (Fig. 1i, u). Taken together, these data suggest rete ridges exhibit a connected 3D network morphology, the bottom of the rete ridges showed enhanced ALP staining, and this 3D rete ridge complex does not regenerate in lesions.

**Figure 1.**
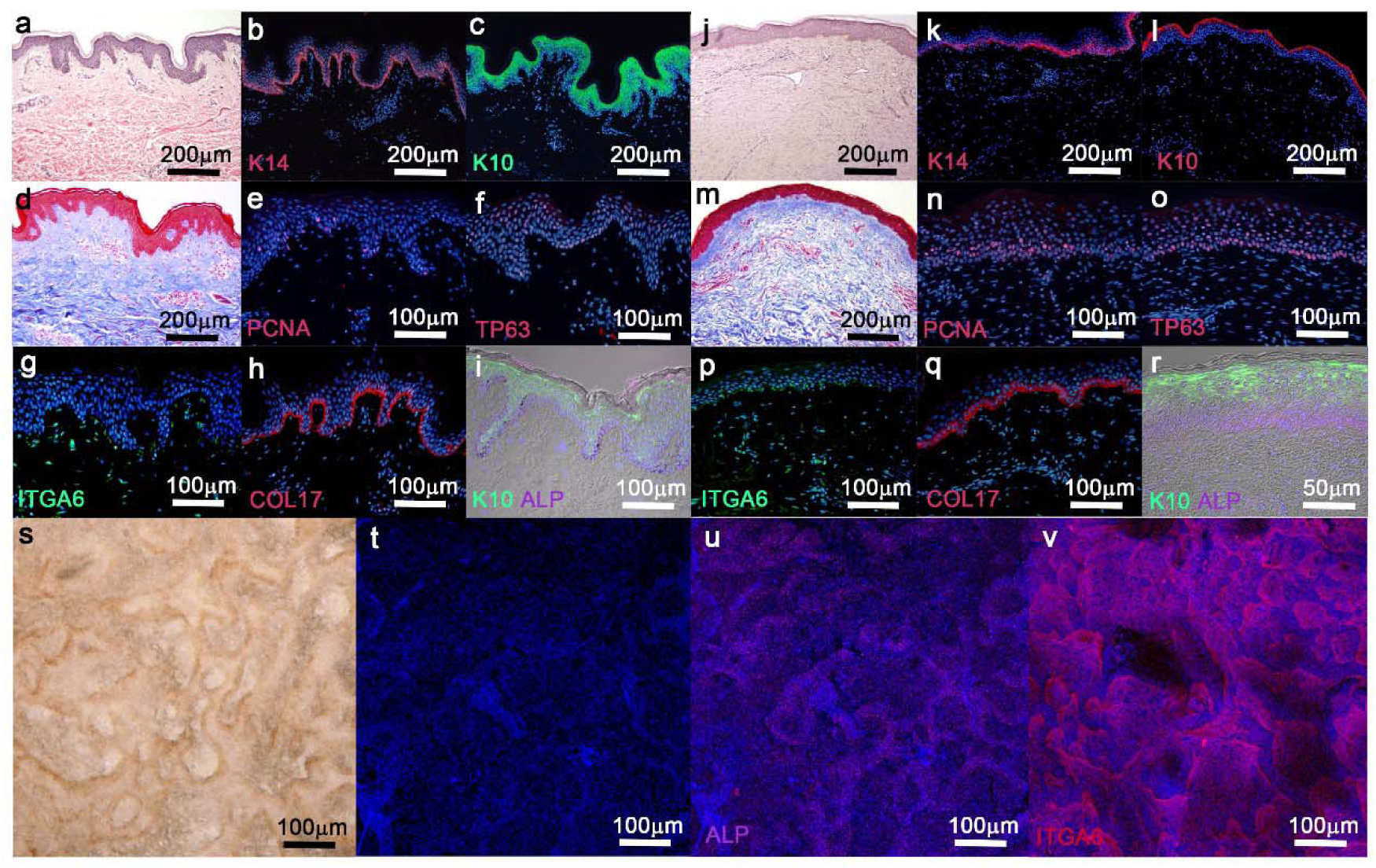
Human rete ridges exhibit 3-dimensional structure and do not regenerate. (a-i) Normal human skin histology and immunohistochemistry; (a) H&E, (b) cytokeratin 14, (c) cytokeratin 10, (d) trichrome, (e) PCNA, (f) TP63, (g) ITGA6, (h) collagen 17, and (l) cytokeratin 10 with alkaline phosphatase stain. (j-r) Healed human skin lesion histology and immunohistochemistry; (j) H&E, (k) cytokeratin 14, (l) cytokeratin 10, (m) trichrome, (n) PCNA, (o) TP63, (p) integrin alpha 6, (q) collagen 17, and (r) cytokeratin 10 with alkaline phosphatase stain. (s-t) Normal human skin was separated and the epithelium viewed from the ventral side; (s) rete ridge network with brown melanin pigment, (t) Hoechst stain, (u) alkaline phosphatase stain, and (v) integrin alpha 6. (blue nuclei; Hoechst)

### Fraser’s Dolphin Gross Wound Healing

Cookie-cutter sharks (*Isistius sp.*) prey upon cetaceans by biting and removing significant sections of LFT skin creating circular wounds approximately 8 cm in diameter (Dwyer, 2011, Grace et al, 2018). Surprisingly, these LFT cetacean wounds heal with little to no scar, unlike human cookie-cutter shark or LFT wounds which scar and require extensive tissue grafts (Honebrink et al, 2011). Of note, an individual cetacean sustains multiple bite wounds, each at a different wound healing time point. Exploiting this, Fraser’s dolphins (FD) LFT skin wound regeneration was characterized (Fig 2a-b and Extended Data Fig 1a-c). The gross healed wound showed a bilateral symmetry that deformed when excising specific wound regions (Fig 2c-e and Extended Data Fig 1d-e). We surmise the wound healed with an inherent tension. This is interesting because tension wounds in humans produce hypertrophic or keloid scars (Hsu et al, 2018). For cetaceans, this mechanism is occurring in a viscous, aqueous, and high shear force environment and previous reports have demonstrated tissue tension modulates stem cell proliferation and differentiation (Ning et al, 2021). Utilizing paraffin sectioning, an overlay tracing of transverse sections parallel to the skin surface was created (Fig 2f, Extended Data Fig 1d-j, and Sup Movies 1-6). Unwounded skin showed an anterior to posterior alignment of rete ridges and included smaller epithelial connections in between agreeing with previous reports (Sokolov, 1955, Stromberg, 1989) (Fig 2f far left or right, Sup Movies 1-6). Interestingly, the healed wound rete ridges were slightly disorganized but were more horizontally aligned (Fig 2f center). This data suggests FD healed LFT wounds with tension and misaligned rete ridges, and this contrasts to humans which scar under tension and do not regenerate rete ridges (Fig 1).

**Figure 2.**
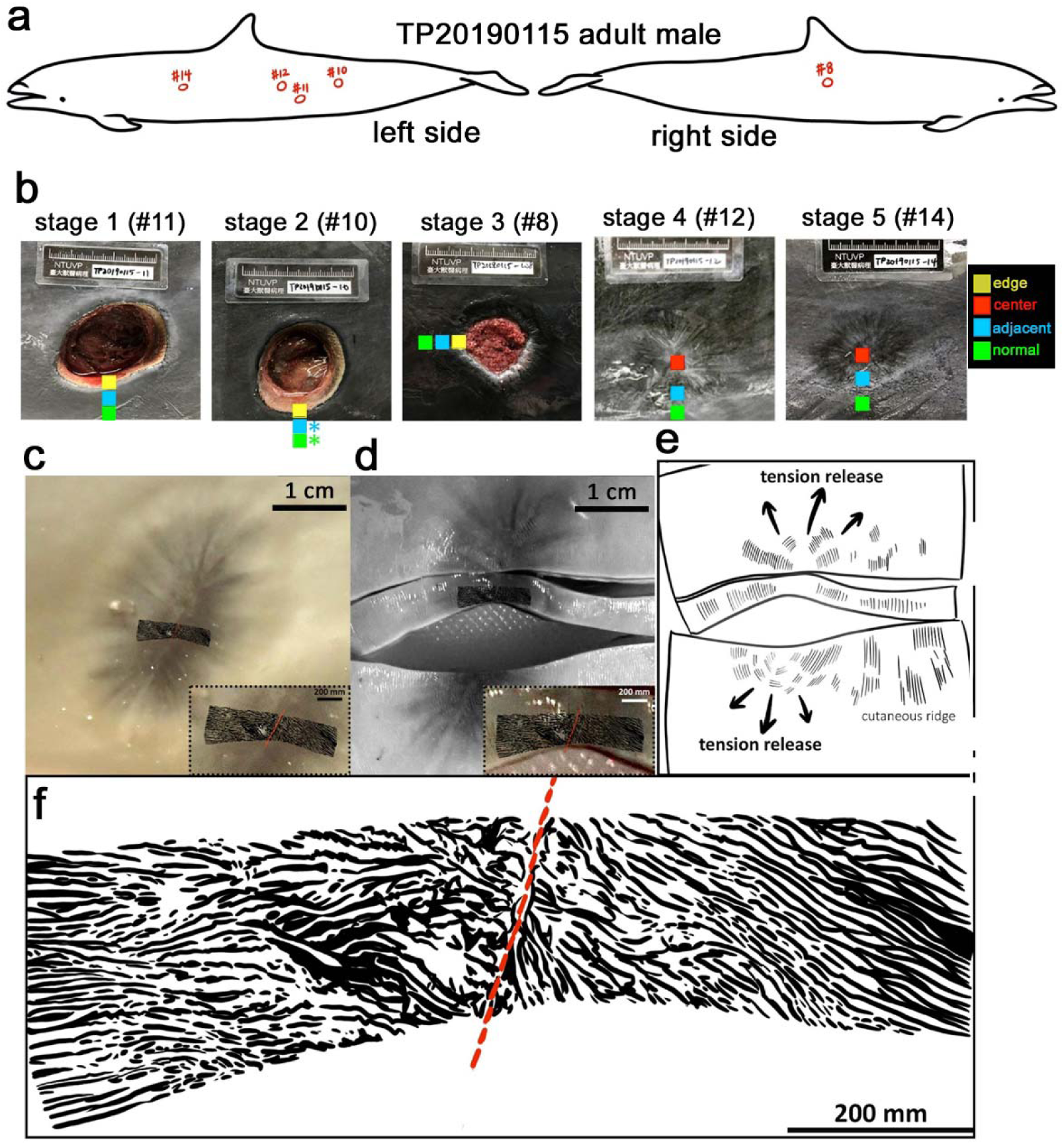
FD cookie-cutter shark bite sites, healing stages, and rete ridge alignment. (a) Wound collection sites. (b) Wound stages; stage-1; new, stage-2; partially contracted, minimal granulation tissue, stage-3; more mature, contracted, granulation tissue, stage-4; reepithelized, pigmentation irregularities, granulation tissue, and stage-5; healed, remodeled, reduced pigmentation irregularities and minimal scar. (yellow box; wound edge, red box; wound center, blue box; wound adjacent, green box; normal skin, blue/green *; tissue collected out of view) (c) Stage-4 before and (d) after dissection. (scale bar; 1 cm, dotted black box; magnified view, scale bar; 200 mm, red line; healed wound mid-line) (e) Tension and cutaneous ridge orientation diagram. (f) Rete ridge overlay tracing for (d). (scale bar; 200 mm, red line; healed wound mid-line)

### *De Novo* Rete Ridge Regeneration

FD rete ridges showed complex morphology with epithelial elongations extending into the papillary dermis (Fig 3 and Extended Data Fig 2). These rete ridges exhibited branches along the lateral and basal sides, and are similar to the ‘knobs or nodules’ reported in Bottlenose dolphins (*Tursiops tuncatus*) (Stromberg, 1989). In comparison, human rete ridges do not extend as deep nor contain branches (Fig 1)(Lin et al, 2019).

**Figure 3.**
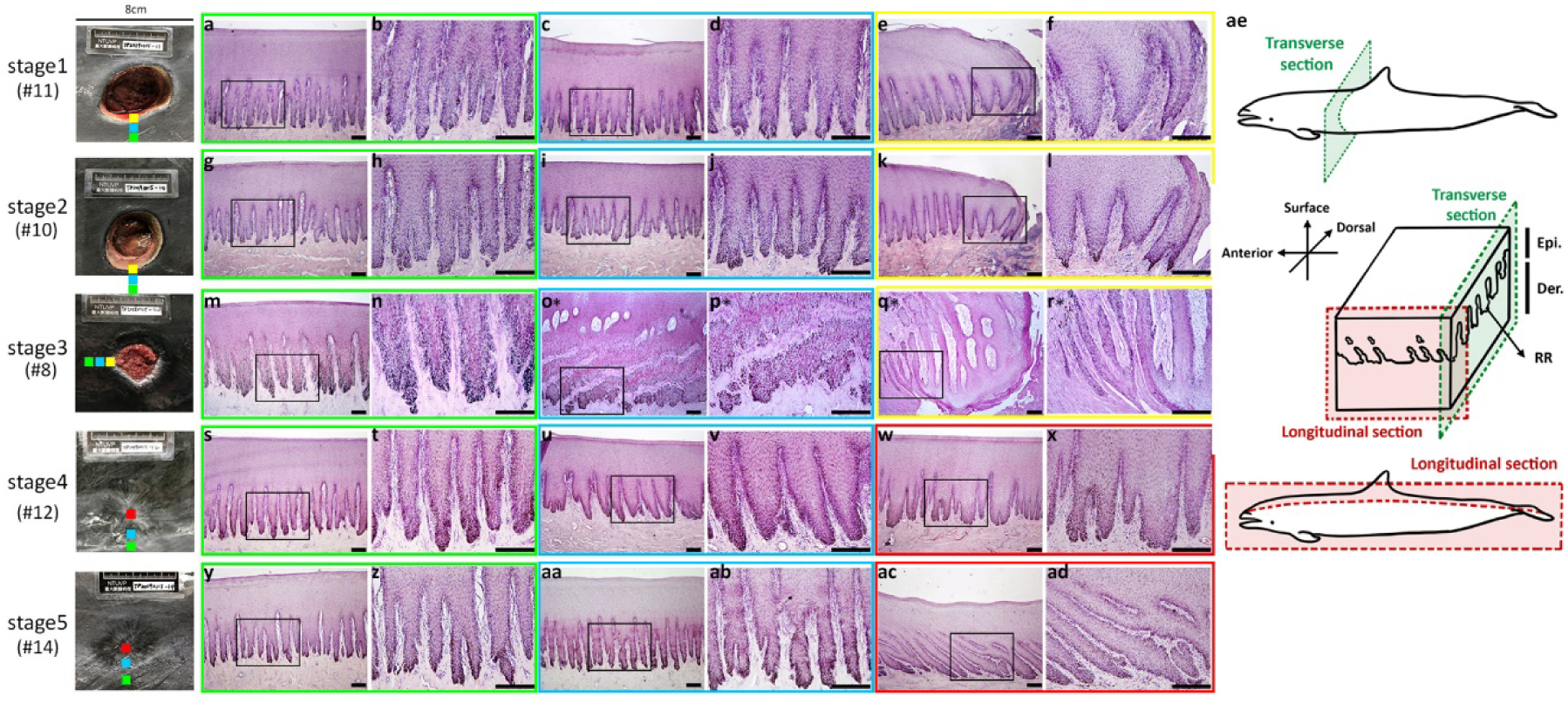
FD rete ridge morphology and de novo regeneration. (a-ad) H&E of unwounded, adjacent, and wound stages; (a-f) stage-1, (g-l) stage-2, (m-r) stage-3, (s-x) stage-4 and (y-ad) stage-5. (a-b, g-h, m-n, s-t, y-z) Unwounded normal rete ridge morphology (green boxes). (c-d, i-j, o*-p*, u-v, aa-ab) Wound adjacent stage-1 (c-d) and 2 (i-j) normal morphology, but stage-3 (o*-p*) and 4 (u-v) abnormal rete ridges. (aa-ab) Stage-5 wound adjacent normal morphology (blue boxes). (e-f) Stage-1 and 2 (k-l) wound edge abnormal rete ridges. (q*-r*) Stage-3 wound edge migrating epithelial tongue with elongating epithelial strands (yellow boxes). (w-x) Stage-4 and (ac-ad) stage-5 wound center and de novo regenerated rete ridges (red boxes). (ae) Animal axes and sectional plane relationship. (scale bars; 200 μm, o*-r*; longitudinal section)

Previously, our group classified cetacean wound healing starting with stage 1 as a recently created wound, and progressing along a continuum towards a completely healed wound at stage 5 (Su et al, 2022). The morphology of unwounded tissues for all stages (1-5) are similar and exhibit healthy normal tissue structure with intact epithelium, basement membrane overlying a papillary dermis (Fig 3). FD stage 1 wound adjacent tissue showed rete ridge structure similar to unwounded tissue (Fig 3a-d and Extended Data Fig 3a-b). Stage 1 wound edges showed a reduced number of rete ridges with increased width and absence of branching (Fig 3e-f, Extended Data Fig 3c, and Extended Data Fig 4). Stage 2 wound edges exhibited similar morphology to stage 1 (Fig 3g-j, Extended Data Fig 3d-e, and Extended Data Fig 4) except epithelial tongue formation at the wound edge (Fig 3k-l and Extended Data Fig 3f). The overlay tracings of healed wounds exhibited rete ridges that were misaligned but more anterior-posterior oriented versus dorsal-ventral (Fig 2f), and we inquired if rete ridges regenerated. Due to the bilateral wound healing symmetry (Fig 2), the stage 3 sample sites were excised in the anterior to posterior plane and sectioned in a longitudinal orientation. Longitudinal sections exhibited a plate-like structure of rete ridges distinct to finger-like structures observed in transverse sections (Fig 3, Extended Data Fig 3, and Extended Data Fig 5).

Stage 3 unwounded tissues exhibited normal morphology (Fig 3m-n, Extended Data Fig 3g-h, and Extended Data Fig 5). The stage 3 wound adjacent tissue exhibited plate-like rete ridge structures obliquely orientated to the wound surface (Fig 3o-p and Extended Data Fig 5). The stage 3 migrating epithelial tongue developed small, thin elongating epithelial strands extending upwards from the basal side at the dermal-epidermal junction (Fig 3q-r, Extended Data Fig 3i, and Extended Data Fig 5). Intriguingly, the rete ridges regenerated *de novo* first, followed by branch formation (Fig 3r, Extended Data Fig 3i, and Extended Data Fig 5). Stage 4 unwounded tissue was normal and wound adjacent tissue demonstrated a reduced number of wide rete ridges (Fig 3s-v, Extended Data Fig 3j-k, Extended Data Fig 4, and Extended Data Fig 5). The stage 4 healed wound center showed rete ridges that were wider and less branched than unwounded tissue (Fig 3w-x, Extended Data Fig 3l, Extended Data Fig 4, and Extended Data Fig 5). Stage 5 unwounded and wound adjacent tissue was unremarkable (Fig 3 y-ab, Extended Data Fig 3m-n, and Extended Data Fig 5). Stage 5 wound center regenerated rete ridges exhibited normal structure with deep epithelial elongations containing branches (Fig 3ac-ad, Extended Data Fig 3o, Extended Data Fig 4, and Extended Data Fig 5). These data suggest FD rete ridges develop complex morphology, regenerate *de novo* first, with branches forming secondarily. LFT human wounds heal with severe scar (Gurtner, Callaghan and Longaker, 2007) and rete ridges do not regenerate (Fig 1).

### Vasculature Regenerates after Rete Ridge

Since humans and FD are both tight-skinned mammals, we examined wound protein expression patterns of basal and suprabasal cell markers for comparison. Stage 1-2 unwounded, wound adjacent, and wound edge tissues showed exclusive expression of cytokeratin 5 (K5) in the basal layer and K17 in the suprabasal layer (Fig 4a-f and Extended Data Fig 6). Importantly, cetacean adaptation utilizes K17 as a suprabasal cell keratin but humans use K10 (Ehrlich et al, 2019). This is interesting because K17 is normally expressed in skin appendages and is activated during wound healing for terrestrial mammals (Zhang, Yin and Zhang, 2019) suggesting cetacean skin is primed or in a state of ‘readiness’ for wound healing. As noted above, the tension of the wound, along with environmental shear forces acting on the wound, could contribute to enhanced activation of skin stem cells creating a synergistic effect with a ‘primed’ state denoted by the K17 biomarker (Ning et al, 2021). At stage 3, the unwounded and wound adjacent tissue expression patterns were similar to stages 1-2 (Fig 4g-h and Extended Data Fig 6). However, the stage 3 migrating epithelial tongue showed expanded K5 expression not restricted to the basal layer in the *de novo* rete ridges, and K17+ cells emerged later during rete ridge maturation (Fig 4i and Extended Data Fig 6). After stage 3, stages 4-5 reverted to normal basal K5 and suprabasal K17 expression patterns (Fig 4j-o and Sup Fig 5). Stage 3 epithelial tongue serial sections showed multiple epithelial strands emanating from a specific epithelial tongue site (Fig 4p-r and Extended Data Fig 6) with co-expression of K5 and K17 near the *de novo* rete ridge tips (Fig 4s-t and Extended Data Fig 6). The rete ridge branching piqued our interest and further analysis of unwounded H&E data showed vascularity in close proximity to branches (Fig 3-4 and Extended Data Fig 2-3, 5). Alpha smooth muscle actin (αSMA) and transgelin/smooth muscle protein 22 alpha (SM22) identify vasculature smooth muscle cells. αSMA and SM22 protein expression patterns identified blood vessels interspersed between rete ridges, confirming previous reports (Menon et al, 2022), and juxtaposed to branches of unwounded tissue (Extended Data Fig 7). We posed the question whether or not *de novo* rete ridge branches regenerated this vascularity association? Vascularity regenerated in between and after the *de novo* rete ridge elongation, but before rete ridge branching (Fig 4u-v and Extended Data Fig 7). Furthermore, the vascularity re-associated with the rete ridge branches (Fig 4v and Extended Data Fig 7w). Previous research documented greater cetacean epidermal keratinocyte proliferation rate insinuating an increased metabolic demand (Hicks et al, 1985). The close association of the rete ridge branches with vascularity could support this metabolic demand by increasing surface contact area and is a current focus for our research group. Taken together, these data suggest FD LFT wounds undergo *de novo* rete ridge regeneration in a specific sequence; rete ridges first, followed by vascularity, and finally branching.

**Figure 4.**
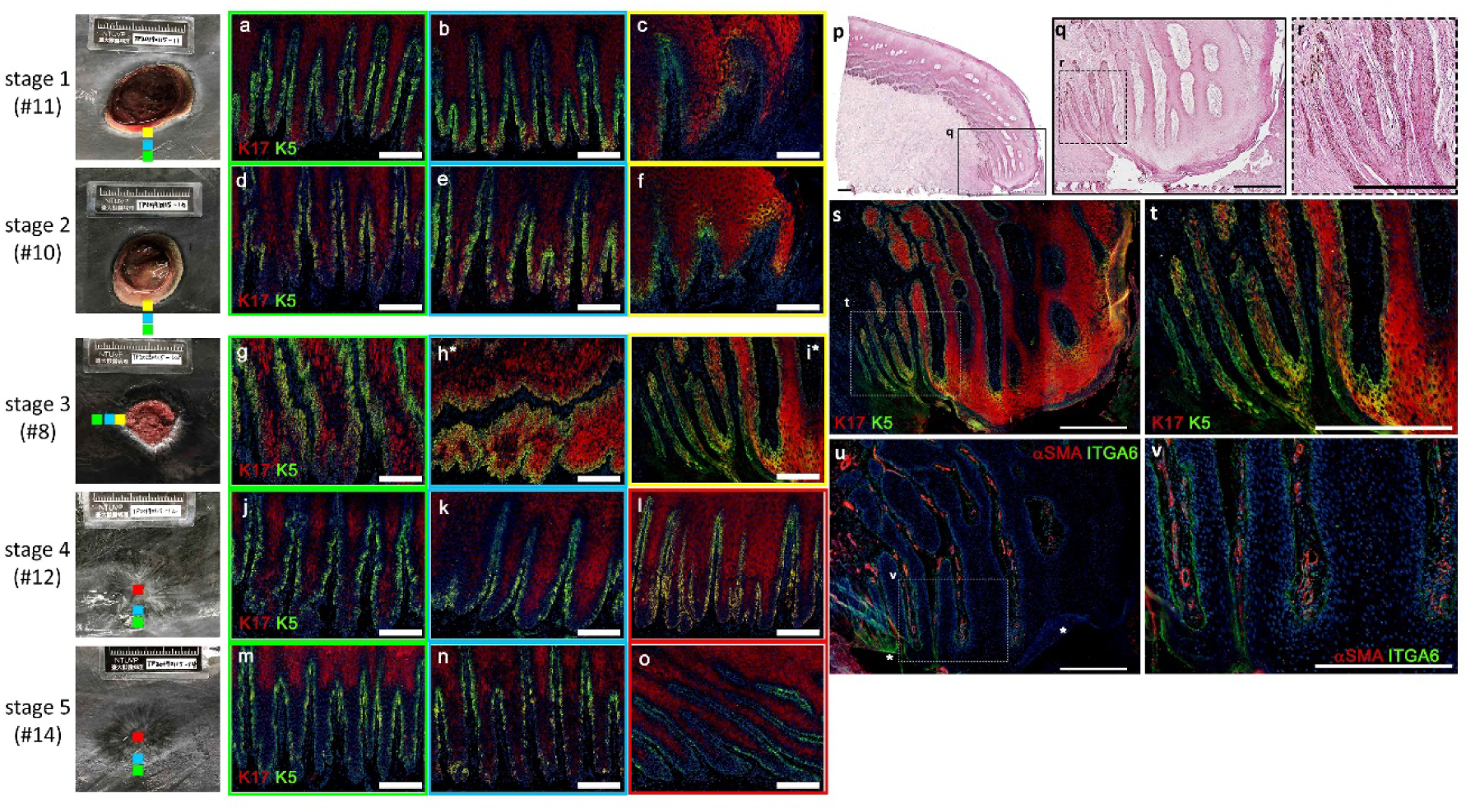
Wounds regenerate complex rete ridge morphology. (a-o) K17 and K5 immunohistochemistry; (a) stage-1 unwounded, (b) wound adjacent, and (c) wound edge; stage-2 (d) unwounded, (e) wound adjacent, and (f) wound edge; stage-3 (g) unwounded, (h*) wound adjacent, and (i*) wound edge; stage-4 (j) unwounded, (k) wound adjacent, and (l) reepithelized wound; stage-5 (m) unwounded, (n) wound adjacent, and (o) reepithelized wound. (scale bars;100 μm, blue; Hoechst, green box; unwounded, blue box; wound adjacent, yellow box; wound edge, red box; wound center, h*-i*; longitudinal section) (p-r) Stage-3 H&E; (p) wound edge, (q) epithelial tongue, and (r) de novo rete ridge formation. (s-v) Stage-3 epithelial tongue immunohistochemistry; (s-t) K17 and K5, or (u-v) α-SMA and ITGA6. (scale bars; 200μm, p-v; longitudinal section, asterisk; tissue fold)

**Figure 5.**
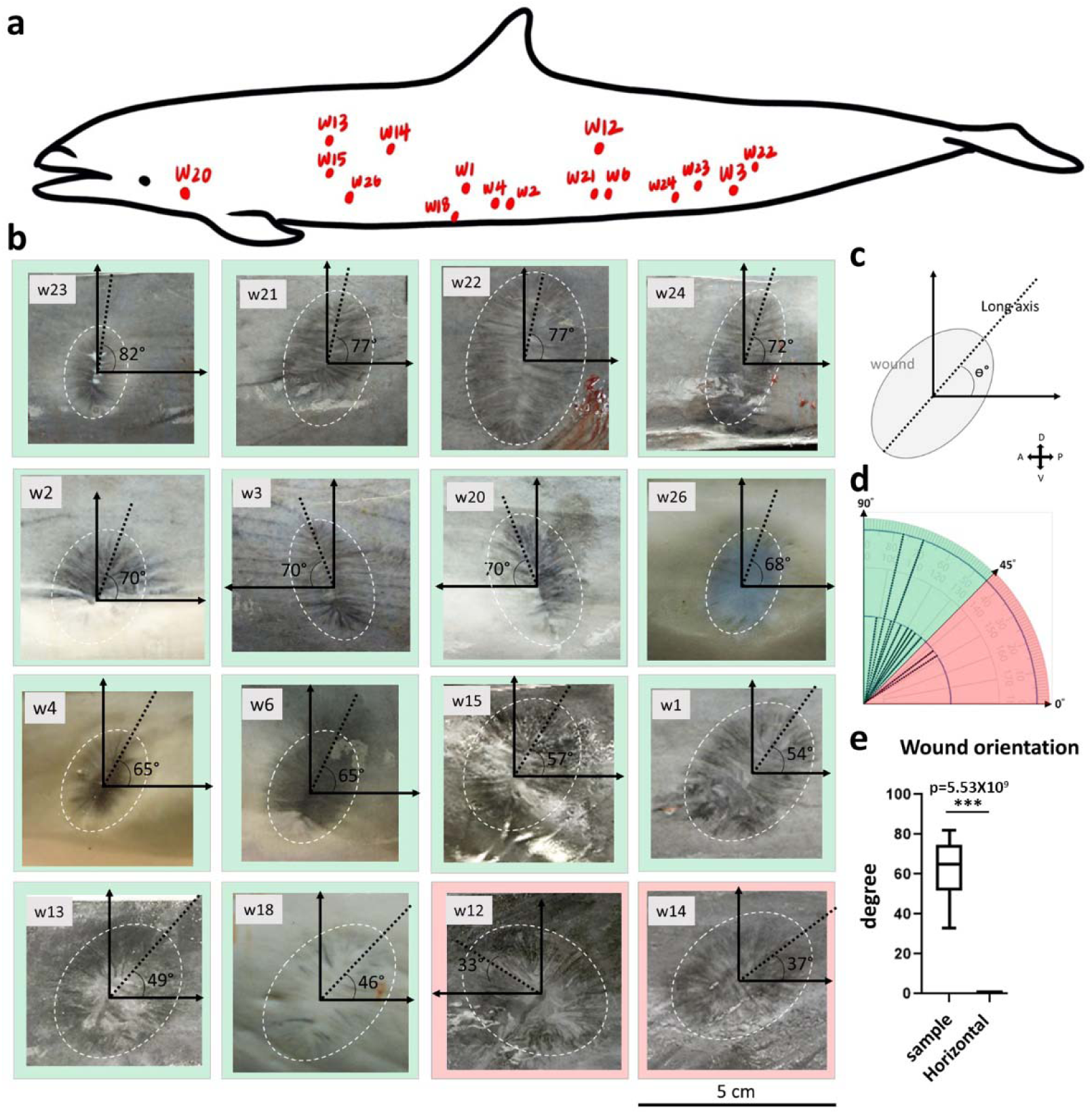
FD bi-lateral wound healing symmetry orientated perpendicular to body axis. (a) Wound collection sites. Adult FD samples #1-3, #12-15, #18, #20-24, #26 from dolphin TP20190115; #4 wound from juvenile dolphin TT20210324; and #6 wound from juvenile dolphin PT20201109. (b) Wound gross pictures showed bi-lateral symmetry with prominent mid-line and radial contraction lines. (white ellipse; wound margin, black dashed line; wound long axis, scale bar; 5 cm) (c) Wound long axis and angle ° to horizontal (A-P) plane measurement. (A; anterior, P; posterior, D; dorsal, V; ventral) (d) Angle distribution histogram from healed wounds. (green color; angle > 45° (n=14 wounds), red color; angle < 45° (n=2 wounds)) (e) Box plot of wound angles with control angle 0°. (one tailed t-test, ***; p<0.001)

### FD LFT Wounds Regenerate in the Anterior-Posterior Axis

Plastic surgeons utilize surgical incision and resulting wound orientation to minimize adverse effects (Amin et al, 2020). FD LFT healed wounds exhibited a bilateral symmetry (Fig 2 and Extended Data Fig 1). We observed *de novo* rete ridge regeneration only in the anterior-posterior orientation and not dorsal-ventral (Fig 3-4 and Sup Fig 3, 5-6). To help elucidate why, multiple healed wounds from multiple FD individuals were analyzed (Fig 5a). The majority of healed wound shapes showed a bilateral symmetrical oval with the long axis of the wound perpendicular to the anterior-posterior plane (Fig 5b-d). Furthermore, the healed wound axes orientated significantly more vertical (dorsal-ventral) than horizontal (anterior-posterior) (n=16, pVal=0.5*10-9) (Fig 5e). These data suggest FD LFT wounds healed with a specific orientation to the body axis and contracted with a bilateral symmetry parallel to the anterior-posterior plane.

## DISCUSSION

Complete, well-organized restoration of mature rete ridge architecture after LFT injury is uncommon in adult tight-skinned terrestrial mammals and typically absent in human scarring. In contrast, here we show Fraser’s dolphin exhibits robust reconstitution of rete/dermal ridges with associated vascular remodeling despite an aquatic, high-shear, high-tension environment.

We characterized the rete ridge regenerative process in LFT wounds of the tight-skinned mammal FD. Of note, human rete ridges show an interconnected reticular network of epithelium with ALP+ basal keratinocytes at the rete ridge bottom that do not regenerate after LFT wounding (Fig. 1). FD LFT wounds heal with minimal defect in an aqueous environment under force (Fig. 2). FD Rete ridges exhibit a longer and branched morphology that regenerates *de novo* with epithelial strands emanating from the migrating epithelial tongue (Fig. 3). The main body of the epithelial rete ridge starts to regenerate, then the vascularity in the papillary dermis begins, followed by branching of rete ridges (Fig. 4). FD LFT wounds contract in the anterior-posterior axis with partial alignment of the regenerated rete ridges (Fig. 1 and 5). Importantly, this regeneration occurs under a relatively higher force and in an aqueous environment versus terrestrial mammal wound healing.

### Contradictory Regeneration

FD healed LFT wounds with little to no noticeable gross tissue deficit (Fig 2-4). This remarkable ability to regenerate LFT wounds is significantly different from the human wound healing result of immense scarring (Gurtner, Callaghan and Longaker, 2007). Counterintuitively, this cetacean skin regeneration occurs in a viscous wet environment (water vs air), and within a stiffer wound architecture. In good clinical practice, human wounds should be kept relatively dry (moist but not wet) to prevent maceration or adverse effects occur (Whitehead et al, 2017). LFT wounds in humans heal with severe scar (Shieh and Cheng, 2015). Additionally, stiff human wounds induce hypertrophic or keloid scar formation with no regeneration of rete ridges or ectodermal organs (Hsu et al, 2018, DeJong et al, 2020). Finally, wound stiffness gradients characterized in mouse LFT wounds showed soft wound areas regenerated while stiff wound areas scarred, and the spiny mouse exhibits enhanced regeneration in part due to a low stiffness wound environment (Harn, H. I. et al, 2021, Harn, Hans I-Chen et al, 2022). How this contradictory regeneration occurs requires further study.

### Regenerated Tissue Architecture

Interestingly, the FD rete ridges regenerated in a similar orientation as the original tissue rete ridge alignment (Fig. 2). Furthermore, the FD wounds contracted more in the anterior-posterior axis versus the dorsal-ventral axis (Fig. 5). This suggests FD wound healing is restoring an anterior to posterior patterning, and contrasts to human LFT wounds that do not regenerate rete ridges, with significant contraction from multiple directional planes. Although partial-thickness human and pig wounds exhibit limited rete ridge regeneration ability the resulting architecture is compromised (Shieh and Cheng, 2015, Lin et al, 2019). Plastic surgeons utilize tension lines to minimize scar formation when making and suturing incisions (Paul, 2018). The FD rete ridges regenerated in the same direction as the contractile force and can be considered regenerating along tension (Fig. 2). Human LFT wounds do not regenerate but induce severe scar under directional force (Hsu et al, 2018). Understanding how FD wound healing promotes regeneration along tension lines could improve human clinical outcomes and is a focus of future studies.

Variable levels of abilities to regenerate adult organs are observed throughout the animal kingdom; zebrafish hearts (Beffagna, 2019), Xenopus tails (Tseng and Levin, 2008, Aztekin et al, 2019, Dunlap and Whited, 2019), salamander limbs (Seifert, Monaghan et al, 2012), opossum digits (Yu et al, 2010), mouse nails (Pulawska-Czub et al, 2021), mouse hair (Hughes et al, 2018), and rabbit hair (Orasan et al, 2016). Loose-skinned animals exhibit remarkable abilities to regenerate skin and hair after LFT wounds. Breedis and Ito characterized *de novo* hair regeneration in rabbit and mice respectively (Breedis, 1954, Ito et al, 2007). Additionally, the wound bed exhibits complex spatiotemporal gene regulation during regeneration (Hughes et al, 2018). Unfortunately, tight-skinned mammals, humans included, lack this ability and severe scar develops.

Partial-thickness wounds demonstrate minimal rete ridge regeneration (Hanson et al, 2016, Lin et al, 2019). How the tight-skinned mammal FD performs *de novo* LFT wound regeneration of complex rete ridge structures exhibiting deep elongated epithelium, epithelial branching, and intimate association with vasculature in a viscous aqueous environment compels further study.

## MATERIALS AND METHODS

### Fraser’s dolphin cookie-cutter shark wounds

Wound samples were collected from four stranded Fraser’s dolphins on the coast of Taiwan; (1) TP20190115 (adult, male, body length: 247 cm, stranded location: Taipei, TW), (2) TT20210324 (juvenile, male, body length: 158.5 cm, stranded location in Taitung, TW), (3) PT20201109 (juvenile, male, body length: 189 cm, stranded location in Pingtung, TW), and (4) IL20191105 (juvenile, male, body length: 221 cm, stranded location in Yilan, TW). Skin sample collecting was coordinated by the groups of; Marine Biology and Cetacean Research Center, National Cheng Kung University, Tainan, Taiwan and Taiwan Cetacean Society. Wound samples were fixed immediately with 10% neutral formalin (Tonyar Biotech. Inc. Taiwan) for TP20190115 or 4% PFA (paraformaldehyde) (Millipore, Billerica MA, USA) for TT20210324 during histological analysis.

### Hematoxylin and eosin staining

Wound tissues were fixed with 10% neutral formalin or 4% PFA overnight at 4°C. Then it was dehydrated in ascending graded ethanol, cleared in xylene, embedded in Paraplast (Sigma, Saint Louis, MO, USA). Tissue sections of 7 μm were affixed to slides, deparaffinized with xylene and rehydrated with ascending ethanol series, and stained with hematoxylin and eosin (Leica, Buffalo Grove, IL, USA) according to accepted protocol. The slides were mounted with Micromount mounting medium (Leica, Buffalo Grove, IL, USA).

### Immunofluorescence staining

The IHC protocol was previously described (Lin et al, 2019). Briefly, tissues sections of 7 μm were affixed to slides, deparaffinized with xylene and rehydrated with descending graded ethanol series. For antigen retrieval, paraffin-embedded sections were incubated in either sodium citrate pH6 or Tris/EDTA, pH 9 at 86°C for 15 min to 1hr depending on tissue condition. The sections were washed and then blocked in 0.6% methanol peroxide at RT (room temperature) for 30 minutes and subsequently incubated with Zeller’s buffer at RT for 1h. Primary antibodies K17 at 1:200 (rabbit polyclonal to cytokeratin 17, Abcam, Cambridge, UK), K5 at 1:200 (Rabbit polyclonal to Cytokeratin 5, Abcam, Cambridge, UK), PCNA at 1:200 (mouse monoclonal, Millipore, Billerica MA, USA), p63 at 1:200 (rabbit monoclonal to p63, Abcam, Cambridge, UK), integrin α6 at 1:200 (rabbit monoclonal to integrin alpha 6, Abcam, Cambridge, UK), α-SMA at 1:200 (mouse monoclonal to alpha-smooth muscle actin, Thermo Fisher Scientific, Eugene OR, USA) and SM22 at 1:200 (rabbit polyclonal to TAGLN/Transgelin, Abcam, Cambridge, UK) were applied on the slides and incubated overnight at 4°C. Next, the slides were washed several times and then fluorescent labeled secondary antibody (Alexa Flour 594 IgG or Alexa Flour 488 IgG Thermo Fisher Scientific, Eugene OR, USA) diluted in Zeller’s buffer added and incubated at 4°C overnight in the dark. Hoechst staining (Abcam, Cambridge, UK) was used for nuclear identification and slides were mounted with glycerin mounting media.

### Serial section movie

Tissues were fixed with 4% PFA overnight at 4°C. Next, dehydrated in ascending graded ethanol, cleared in xylene, embedded in Paralast (Sigma, Saint Louis, MO, USA). Pictures were taken of Fraser’s dolphin wound tissues block-face after each 7 μm slice of the block from the microtome. The pictures were joined together as a series to demonstrate rete ridge orientation at each depth.

### Image acquisition

Gross images were recorded with Olympus TG6. Microscopic images were recorded with BX51 Olympus Fluorescent Microscope. Serial paraffin block-face images were recorded with an Olympus OM-D camera mounted on the photo port of a Olympus SZ61 stereomicroscope. Serial block-face movies were post-processed with ImageJ software (Bethesda, MD, USA). The entire field of view of individual pictures were adjusted for exposure and composite pictures assembled in Adobe Photoshop (San Jose, CA, USA).

### Statistical analysis

All data figures are representative of at least three samples (n≥3). Box plot of the mean and standard deviations were calculated. Unpaired 1 tailed t-test and 1-way ANOVA were performed on data sets to establish significant differences. *; p-value <0.05, **; p-value <0.01, ***; p-value <0.001.

## Supporting information

Movie 1 Normal Skin

Movie 2 Unwounded Skin Anterior to Wound

Movie 3 Wound Adjacent Skin Anterior to Wound

Movie 4 Wound Center

Movie 5 Wound Adjacent Skin Posterior to Wound

Movie 6 Unwounded Posterior to Wound

## ACKNOWLEDGEMENT

We would like to thank the Marine Biology and Cetacean Research Center in National Cheng Kung University and Taiwan Cetacean Society for assistance with sample collection. We also thank the International Research Center for Wound Repair and Regeneration of the College of Medicine at National Cheng Kung University for their wound tissue expertise and critical suggestions for this project. This work is supported by National Science and Technology Council Grants 114-2314-B-006-100, 110-2314-B-006-109, 111-110-2314-B-006-52, 112-2314-B-006-073, The Featured Areas Research Center Program within the framework of the Higher Education Sprout Project by the Ministry of Education, Taiwan, and The Excellent Research Center Program by the Ministry of Science and Technology 108-3017-F-006-002.

## AUTHOR CONTRIBUTIONS

Conceptualization, M.W.H., H.-V.W., W.-C.Y., C.-M.C., T.-Y.L.; methodology, M.W.H., T.-Y.L., H.-V.W.,W.-C.Y., Y.-T. C, T.-H. C; investigation, T.-Y.L., H.-V.W., C.-Y.S., C.-C.Y., S.-J.S., Y.-T. C; writing original draft preparation, M.W.H., T.-Y.L., H.-V.W.; writing, review, and editing, M.W.H., T.-Y.L., C.-M.C., H.-V.W.; supervision, M.W.H., C.-M.C., M.-J.T.; project administration, M.W.H., H.-V.W.,W.-C.Y.; funding acquisition, M.W.H, H.-V.W.. All authors have read and agreed to the published version of the manuscript.

## COMPETING INTEREST DECLARATION

The authors declare no competing interests.

**Correspondence and requests for materials should be addressed to M.W.H.**

---

**Extended Data Figure 1.**
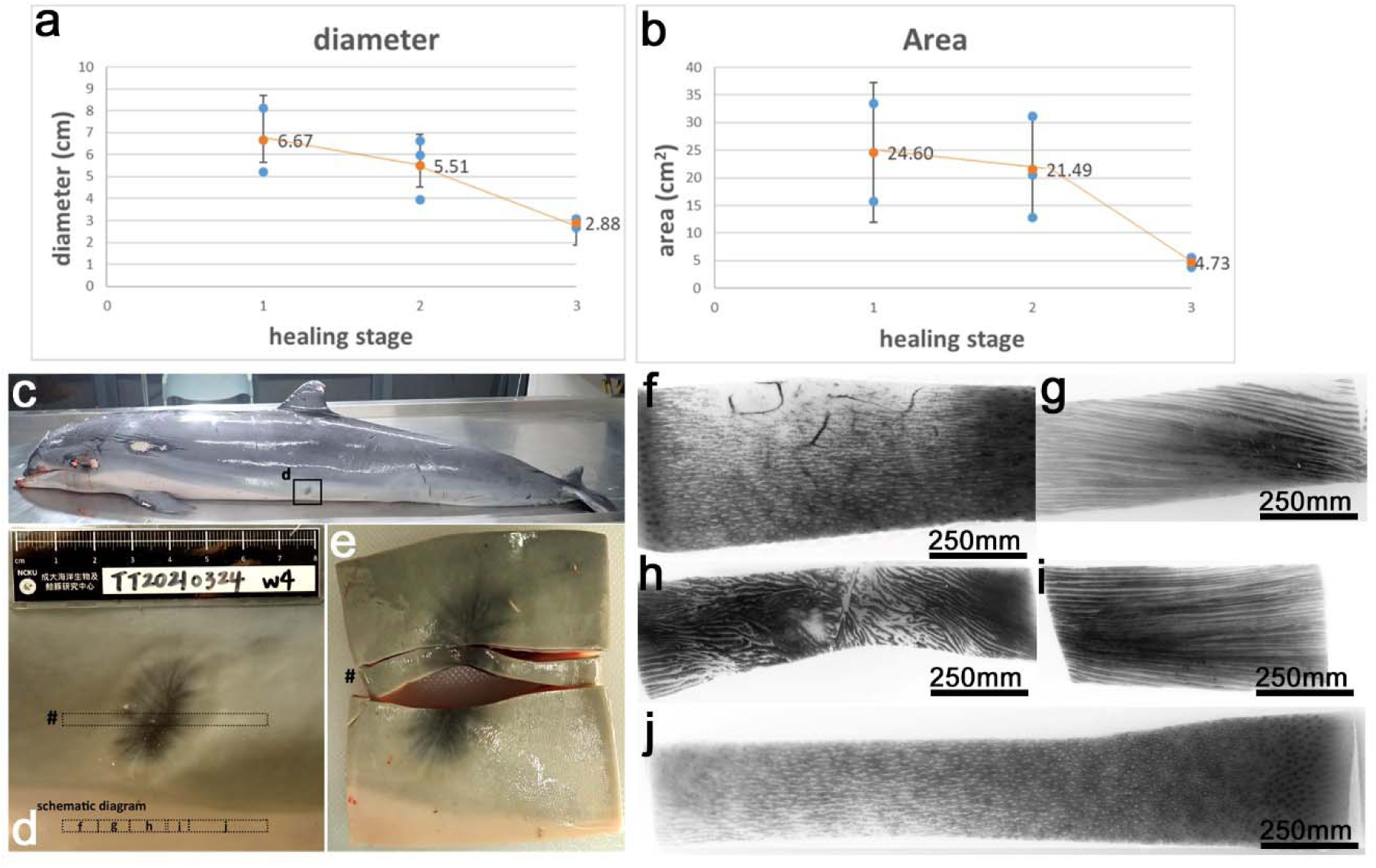
Fraser’s Dolphin wound stage contraction and rete ridge overlay tracing. Wound diameters (a) and areas (b) (mean ± SD) were quantified by calculating image pixels per square centimeter. (n=2 for stage1, n=3 for stage2, and n=2 for stage3) (c) Stage 4 cookie-cutter bite wound was collected from the ventral-lateral side of a juvenile Fraser’s dolphin (TT20210324). (d) Gross picture of stage 4 wound from (c) showed a dorsal-ventral black midline through the center representing the wound long axis with dark contraction lines radiating out. This dissected wound was divided into five sections (black boxes f, g, h, i, j). (#; site of dissection, black box; outline of dissection) (e) Gross picture of dissected stage 4 wound showed deformation due to tension release after cutting. (f-j) Photographs of dissected wound tissue divisions (f, g, h, i, j) embedded into paraffin blocks, sectioned parallel to the skin surface, and used to create the rete ridge schematic overlay diagram for Figure 1f. (f) Unwounded skin, left (anterior) side of the wound. (g) Wound adjacent, left (anterior) side of the wound. (h) Wound center. (i) Wound adjacent, right (posterior) side of the wound. (j) Unwounded skin, right (posterior) side of the wound.

**Extended Data Figure 2.**
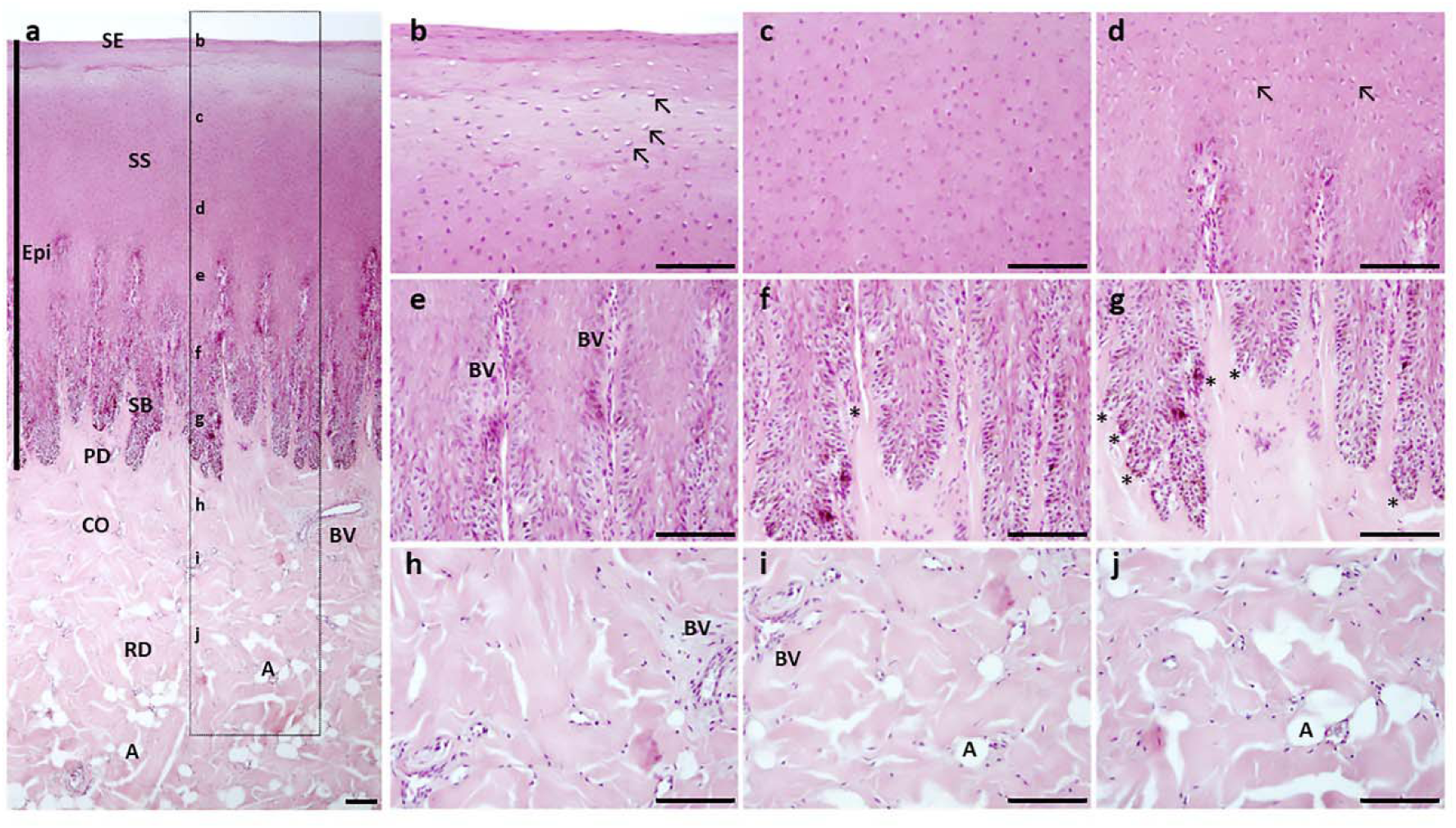
Normal skin architecture of Fraser’s dolphin. (a) H&E staining of normal skin shows a thick epidermis. Epithelium (Epi) of Fraser’s dolphin contains three layers: stratum externum (SE), stratum spinosum (SS), and stratum basale (SB). Prominent elongated rete ridges with small branches extending from the basal and lateral sides are interdigitated with blood vessels in the papillary dermis. Papillary dermis (PD) interspersed and below the rete ridges contains extracellular matrix and vasculature. Reticular dermis (RD) contains blood vessels (BV), collagen (CO) and adipocytes (A). (b-j) Magnified panels from (a) show; (b) cells in the SE with flattened nuclei and the superficial SS demonstrate clear perinuclear cytoplasmic halo and marginated nuclei (black arrow). (c) The mid regional layer of the stratum spinosum shows round nucleated keratinocytes. (d) The keratinocytes of the lower spinosum layer and the top of rete ridges showed elongated morphology with a small population showing clear perinuclear cytoplasmic halos and marginated nuclei (black arrows). (e) Rete ridges interdigitated with blood vessels. (f-g) Rete ridge branches are present at the rete ridge bottom (*). (h) Papillary dermis contains collagen and vasculature. (i-j) Reticular dermis contains collagen, vasculature, and scattered adipocytes (scale bar:100 μm).

**Extended Data Figure 3.**
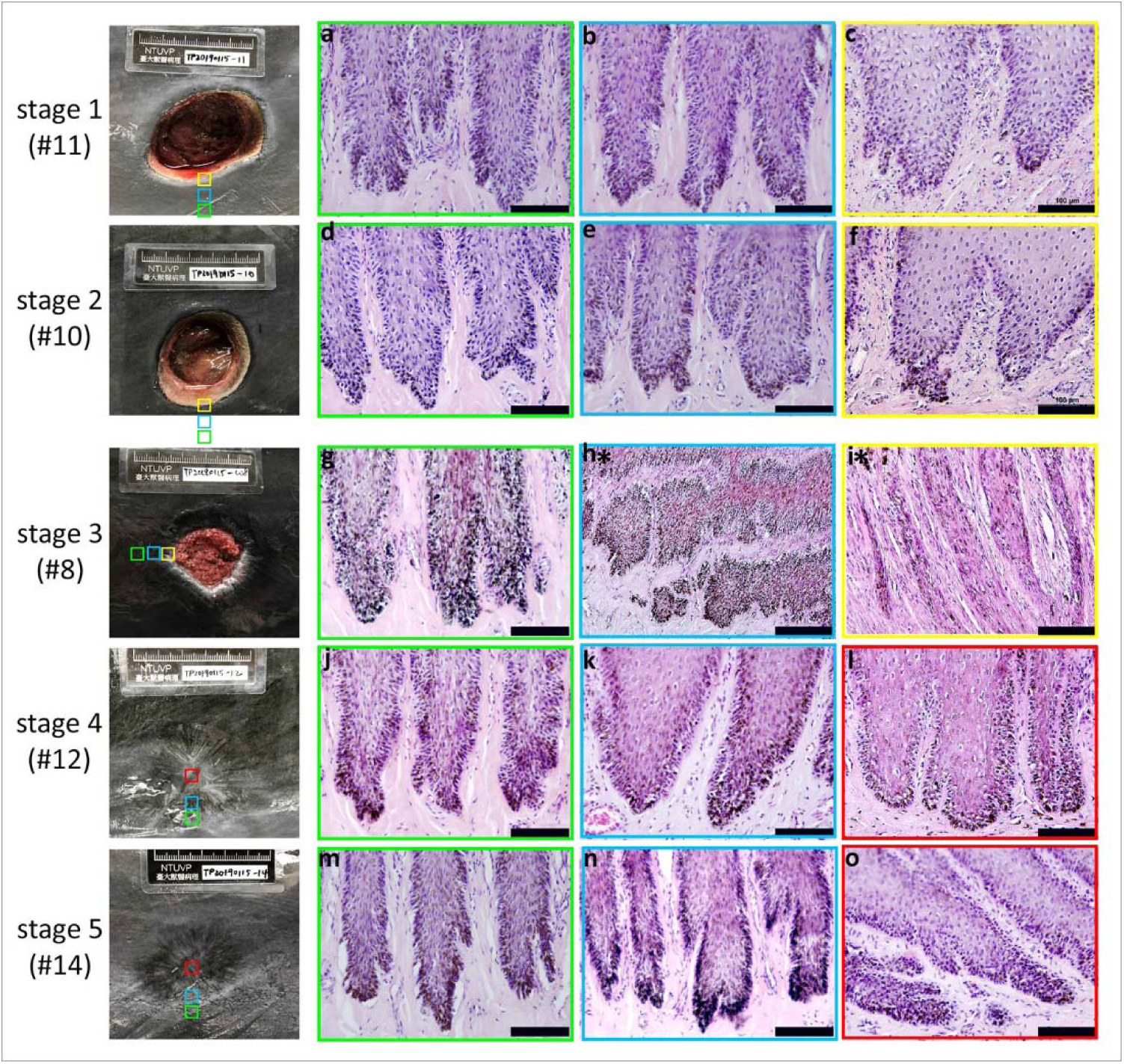
*De novo* rete ridge branch regeneration formed later. (a-o) H&E transverse paraffin sections of Fraser’s dolphin unwounded, wound adjacent, and wound tissues for each healing stage. (a, d, g, j, m) Unwounded skin showed profuse rete ridge branching on the bottom and lateral sides (green boxes). (b, e, h*, k, n) Wound adjacent tissue for stage 1 (b), stage (2) (e), stage 3 (h*, longitudinal section), and stage 4 (k) showed reduced branching of rete ridges (blue boxes). Stage 5 wound adjacent (n) showed normal branching of regenerated rete ridges. (c, f, i*, l, o) Wound edge tissue for stage 1 (c) and 2 (f) showed minimal or non-branched short and wide rete ridges. Stage 3 wound edge tissue (i*, longitudinal section) showed migrating epithelial tongue with elongated epithelial strands without branching (yellow boxes). Stage 4 wound center tissue (l) showed branched short and wide regenerated rete ridges. Stage 5 wound center tissue (o) showed normal sized and profusely branched regenerated rete ridges (n) (red boxes). (scale bars; 100 μm, green box; normal skin, blue box; wound adjacent, yellow box; wound edge, red box; wound center, *h-i*; longitudinal section)

**Extended Data Figure 4.**
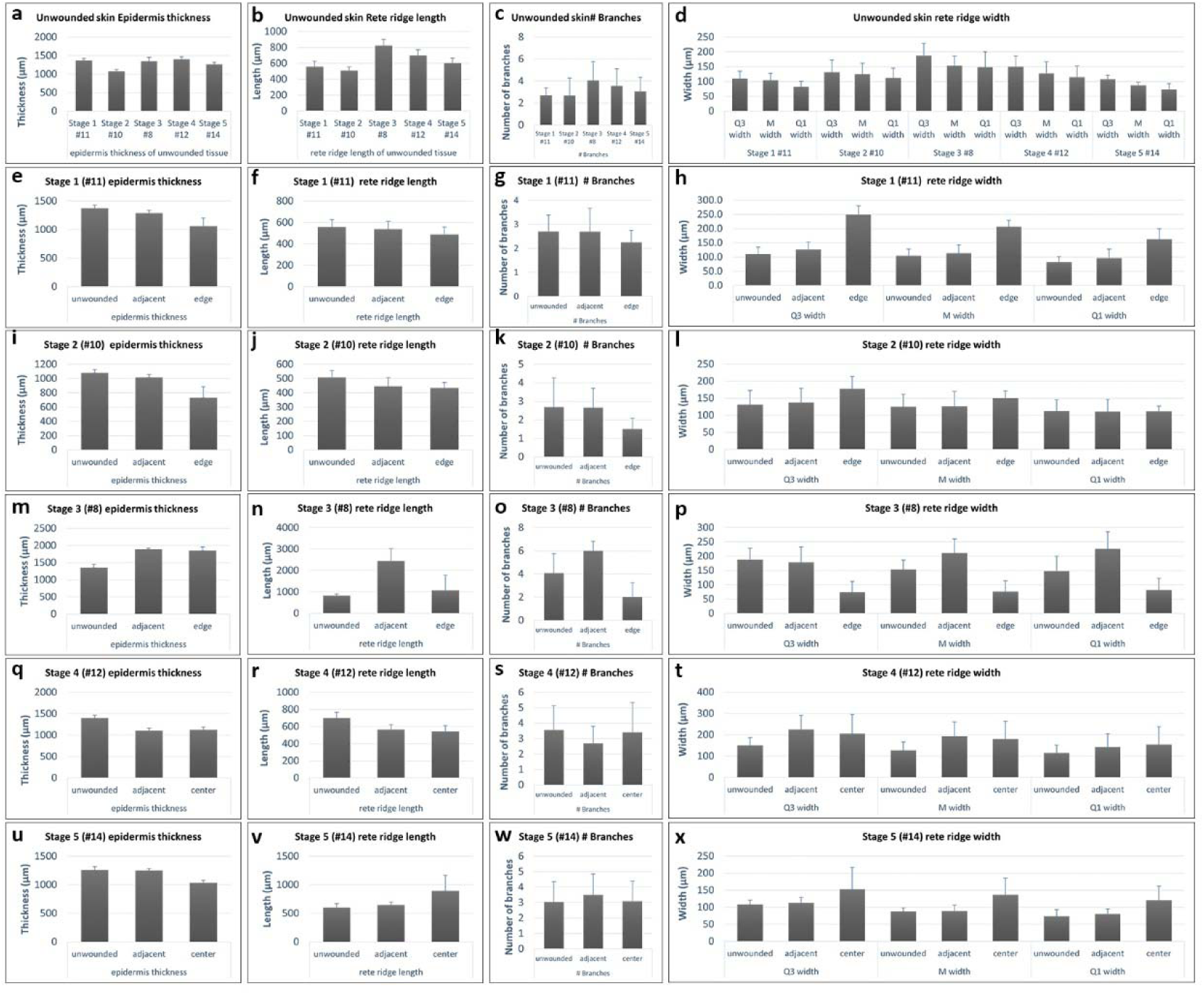
Fraser’s dolphin epithelium thickness, rete ridge length, number of rete ridge branches, and rete ridge width during large full-thickness wound healing. (a-x) Rete ridge physical measurements from unwounded, stage1, stage 2, stage 3, stage 4, and stage 5 wounds for epidermis thickness, rete ridge length, rete ridge branches, and rete ridge width. Rete ridge physical measurement for (a-d) unwounded, (e-h) stage 1 wound (#11), (i-l) stage 2 wound (#10), (m-p) stage 3 wound (#8), (q-t) stage 4 wound (#12), and (u-x) stage 5 wound (#14) epidermis thickness, rete ridge lengths, rete ridges branches, and rete ridge widths. (ANOVA for a-d; Unpaired t-test for e-x; significant difference * p<0.05, ** p<0.01, *** p<0.001, **** p<0.0001)

**Extended Data Figure 5.**
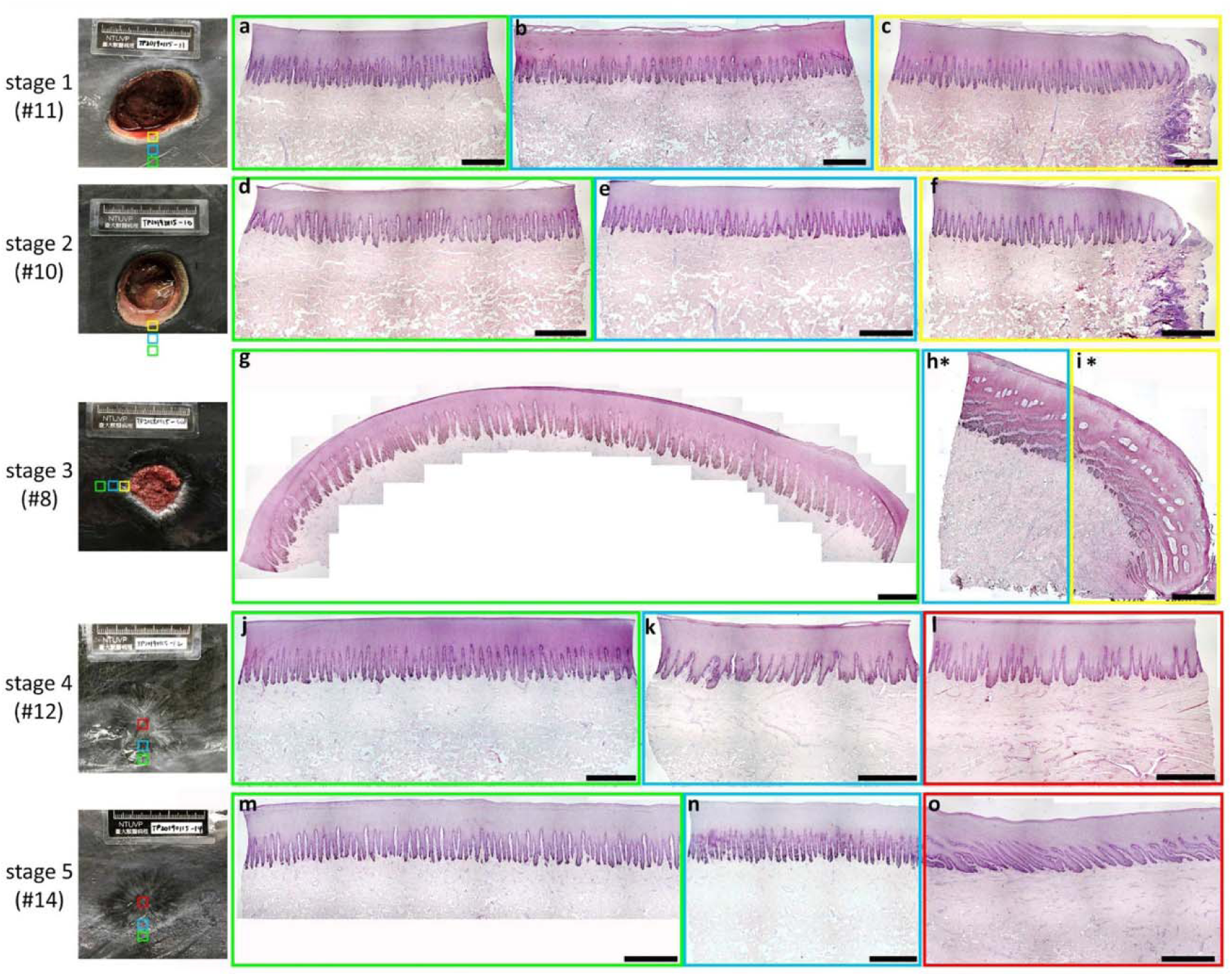
Time course of large full-thickness wound healing and rete ridge regeneration morphology for Fraser’s dolphin cookie-cutter shark bite wounds. (a-o) Global wound view of H&E paraffin sections for Fraser’s dolphin unwounded (green box), wound adjacent (blue box), and wound tissues (yellow box; wound edge or red box; healed wound center) for each healing stage. (a, b, g, j, m) Unwounded skin showed regular, elongated and profusely branched rete ridges for all stages. (b, e, h*, k, n) Wound adjacent tissue for stage 1 (b) and stage 2 (e) showed elongated and branched rete ridges. Stage 3 wound adjacent longitudinal section (h*) showed rete ridge plates with multiple mesenchymal tissue fenestrations. Stage 4 wound adjacent tissue (k) showed regenerated widened rete ridges with reduced branching. Stage 5 wound adjacent (n) showed regular, elongated and branched rete ridges similar in size and structure to unwounded tissue (m). (c, f, i*, l, o) Wound edge tissue for stage 1 (c) and 2 (f) showed short, wide and reduced branched rete ridges. Longitudinal section of stage 3 wound edge tissue (i*) showed an epithelial tongue with elongating strands of epithelium extending from the stratum basale to the stratum spinosum suprabasal layer with mesenchymal fenestrations. The reepithelized stage 4 wound center tissue (l) showed short, wide, and branched regenerated rete ridges. Stage 5 wound center tissue (o) showed long, thin, and branched rete ridges. (scale bars: 1 mm, green box; unwounded skin, blue box; wound adjacent, yellow box; wound edge, red box; healed wound center, *h-*i; longitudinal section)

**Extended Data Figure 6.**
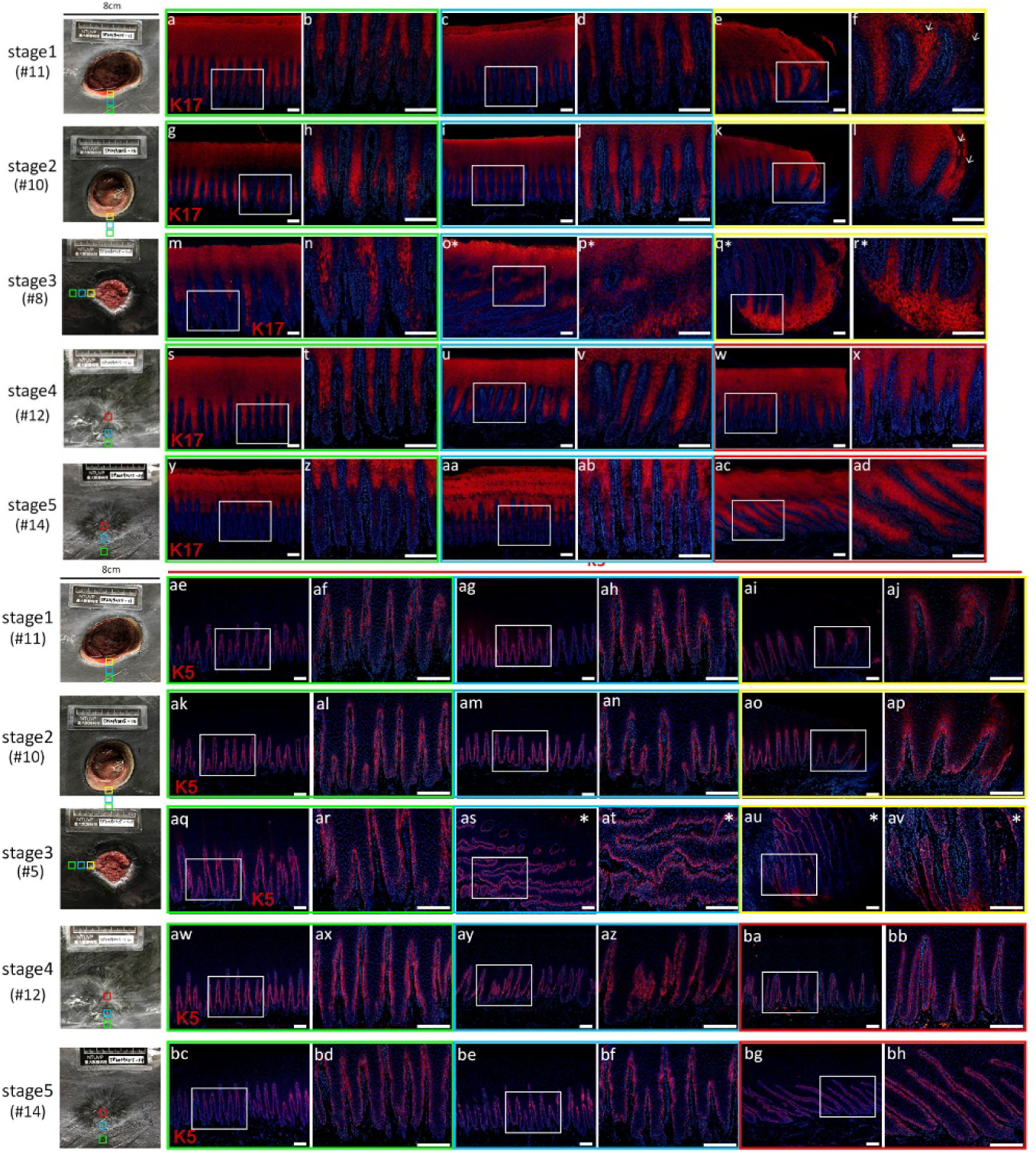
Rete ridge cytokeratin 17 and 5 expression patterns during large full-thickness wound regeneration. Fraser’s dolphin unwounded, wound adjacent, and wound tissue paraffin sections were immunostained with K17 or K5 for all healing stages. (a-f) Stage 1 unwounded (a-b), wound adjacent (c-d), and wound edge (e-f) tissue, (g-l) stage 2 unwounded (g-h), wound adjacent (i-j), and wound edge (k-l) tissue, (m-r) stage 3 unwounded (m-n), wound adjacent (o*-p*), and wound edge (q*-r*) tissue K17 protein expression patterns. (s-x) stage 4 unwounded (s-t), wound adjacent (uv), and reepithelized wound (w-x) tissue, (y-ad) stage 5 unwounded (y-z), wound adjacent (aa-ab), and reepithelized wound (ac-ad) tissue K17 protein expression patterns. (ae-aj) Stage 1 unwounded (ae-af), wound adjacent (ag-ah), and wound edge (ai-aj) tissue, (ak-ap) stage 2 unwounded (ak-al), wound adjacent (am-an), and wound edge (ao-ap) tissue, (aq-av) stage 3 unwounded (aq-ar), wound adjacent (as*-at*), and wound edge (au*-av*) tissue, (aw-bb) stage 4 unwounded (aw-ax), wound adjacent (ay-az), and reepithelized wound (ba-bb) tissue, (bc-bh) stage 5 unwounded (bc-bc), wound adjacent (be-bf), and reepithelized wound (bg-bh) tissue K5 protein expression patterns. (scale bars; 200 μm, red; K17 or K5, blue; Hoechst, green box; normal skin, blue box; wound adjacent, yellow box; wound edge, red box; wound center, o*-r* and as*-av*; longitudinal section; white arrow; missing tissue)

**Extended Data Figure 7.**
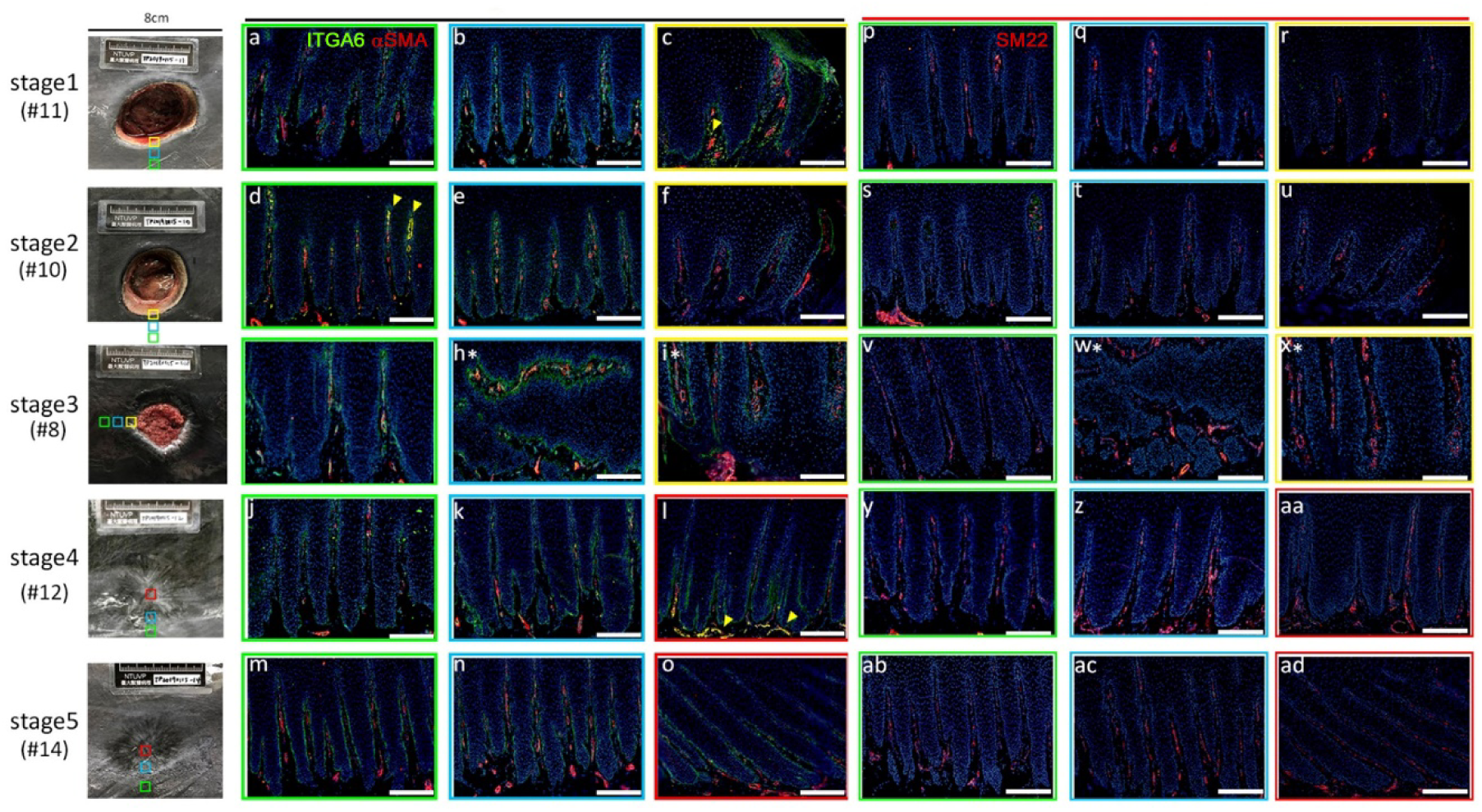
Regeneration of rete ridge associated vasculature. (a-ad) Fraser’s dolphin unwounded, wound adjacent, and wound tissue paraffin sections were immunostained with ITGA6, αSMA, or SM22 for each stage. (a-c, p-r) Stage 1 unwounded (a, p), wound adjacent (b, q), and wound edge (b, r) tissue, (d-f, s-u) stage 2 unwounded (d, s), wound adjacent (e, t), and wound edge (f, u) tissue, (g-i, v-x) stage 3 unwounded (g, v), wound adjacent (h, w), and wound edge (i, x) tissue, (j-l, y-aa) stage 4 unwounded (j, y), wound adjacent (k, z), and reepithelized wound (l, aa) tissue, (m-o, ab-ad) stage 5 unwounded (m, ab), wound adjacent (n, ac), and reepithelized wound (o, ad) tissue ITGA6, αSMA, and SM22 protein expression patterns. (scale bars; 200 μm, green ITGA6, red; αSMA or SM22, blue; Hoechst, green box; normal skin, blue box; wound adjacent, yellow box; wound edge, red box; wound center, h*-i* and w*-x*; longitudinal section, yellow arrowhead; red blood cell autofluorescence)

